# A trimeric human angiotensin-converting enzyme 2 as an anti-SARS-CoV-2 agent in vitro

**DOI:** 10.1101/2020.09.18.301952

**Authors:** Tianshu Xiao, Jianming Lu, Jun Zhang, Rebecca I. Johnson, Lindsay G.A. McKay, Nadia Storm, Christy L. Lavine, Hanqin Peng, Yongfei Cai, Sophia Rits-Volloch, Shen Lu, Brian D. Quinlan, Michael Farzan, Michael S. Seaman, Anthony Griffiths, Bing Chen

## Abstract

Effective intervention strategies are urgently needed to control the COVID-19 pandemic. Human angiotensin-converting enzyme 2 (ACE2) is a carboxypeptidase that forms a dimer and serves as the cellular receptor for SARS-CoV-2. It is also a key negative regulator of the renin-angiotensin system (RAS), conserved in mammals, which modulates vascular functions. We report here the properties of a trimeric ACE2 variant, created by a structure-based approach, with binding affinity of ~60 pM for the spike (S) protein of SARS-CoV-2, while preserving the wildtype peptidase activity as well as the ability to block activation of angiotensin II receptor type 1 in the RAS. Moreover, the engineered ACE2 potently inhibits infection of SARS-CoV-2 in cell culture. These results suggest that engineered, trimeric ACE2 may be a promising anti-SARS-CoV-2 agent for treating COVID-19.

## Introduction

The current COVID-19 pandemic, caused by severe acute respiratory syndrome coronavirus 2 (SARS-CoV-2), has infected more than 29 million people worldwide, leading to over 900 thousand deaths, with devastating socio-economic impacts. Effective intervention strategies are urgently needed to control the pandemic.

Since the outbreak of the virus, several therapeutic approaches have been evaluated in the hope of providing a viable treatment for COVID-19. First, convalescent sera from individuals recovered from the infection were used with encouraging results^1–3^, but also some drawbacks (e.g., batch variations, possible blood-borne pathogens and blood-type matching). Second, patient-derived, potently neutralizing monoclonal antibodies have been isolated, which could provide a more powerful passive immunotherapy than convalescent sera^4–7^. Third, structure-guided design of peptide-based viral entry inhibitors has yielded promising leads in *in vitro* assays^8,9^; their efficacy requires further clinical evaluation. Fourth, known drugs or drug candidates, including remdesivir, favipiravir and ribavirin (viral RNA polymerase inhibitors); lopinavir and ritonavir (viral protease inhibitors); as well as hydroxychloroquine, corticosteroids and interferons (with more complicated antiviral mechanisms), have been repurposed as COVID-19 therapuetics^10,11^. Among them, remdesivir has received Emergency Use Authorizations (EUA) from the U.S. Food and Drug Administration (FDA), while hydroxychloroquine has been shown to be ineffective^12^. Finally, many new therapeutic candidates are in various stages of development (ref^13,14^; https://www.bio.org/policy/human-health/vaccines-biodefense/coronavirus/pipeline-tracker). While the infection resolves on its own in most asymptomatic and mild cases over time, COVID-19 in severe cases appears to progress in two phases – initial active viral replication in the respiratory system and subsequent excessive immune responses leading to multiple organ failure and possible death^15^. Thus, antivirals alone may be insufficient to change the course of disease progression for the population that needs intervention the most if administrated too late.

Human angiotensin-converting enzyme 2 (ACE2) is the cellular receptor for SARS-CoV-2 and binds the receptor binding domain (RBD) of the spike (S) protein of the virus to promote viral entry into the host cells and initiate infection^16,17^. It is a type I membrane glycoprotein containing an extracellular ectodomain that has metallopeptidase activity. Its neck domain near the transmembrane anchor mediates dimerization^18^. ACE2 is also a key negative regulator of the renin-angiotensin system (RAS) - a major hormone system, conserved in mammals and some other vertebrate animals, for modulating vascular function^19,20^. The RAS controls extracellular fluid volume and blood pressure homeostasis by regulating the levels of renin and angiotensins in the circulation. Renin cleaves angiotensinogen to release angiotensin I (Ang I), which can be further processed by angiotensin-converting enzyme (ACE) into angiotensin II (Ang II) - a vasoconstrictive peptide that raises blood pressure and increases the extracellular fluid volume in the body by activating the angiotensin II receptors, including angiotensin II receptor type I (AT1R)^21^. ACE2 primarily converts Ang II to angiotensin-(1–7) (Ang 1–7), which is a vasodilator, thereby counter-balancing the effect of ACE/Ang II and playing critical roles in preventing hypertension and tissue damages^22^.

The protective roles of ACE2 in acute respiratory distress syndrome (ARDS) and acute lung injury (ALI) have been demonstrated in animal models^23–25^. A recombinant soluble human ACE2 (rhACE2) was recently reported to block SARS-CoV-2 infection in cell culture and human organoids^26^, prompting a phase 2 clinical trial for use of rhACE2 as a treatment for COVID-19 patients (NCT04335136). Thus, administration of exogenous ACE2 may be a promising therapeutic strategy for treating COVID-19, because it could not only block viral spread but also modulate the RAS to prevent organ injury. We therefore set out to design a series of ACE2 variants to enhance their binding affinity for SARS-CoV-2 S protein and their potency in blocking SARS-CoV-2 infection.

## Results

### Structures of soluble ACE2 in complex with SARS-CoV-2 S protein trimer

To facilitate design of ACE2-based viral fusion inhibitors, we first determined, by cryo-EM, the structures of a monomeric soluble ACE2 (residue 18-615) in complex with a stabilized soluble SARS-CoV-2 S protein trimer (Fig. S1; ref^27^). We prepared the complex by mixing the two proteins because the monomeric ACE2 dissociates from S trimer very rapidly^28^. After 3D classification (Fig. S2 and S3), we found four distinct classes that represent ACE2-free S trimer in the one-RBD-up conformation, one ACE2 bound S trimer, two ACE2 bound S trimer, and three ACE2 bound S trimer, respectively (Fig. 1). Consistent with previous findings with ACE2 binding to SARS-CoV S protein as well as a recent SARS-CoV-2 study^29^, ACE2 interacts with the RBD in its up conformation. While the NTD (N-terminal domain) of S1 shifts outwards slightly, the S2 portion remains largely unchanged upon ACE2 binding, even when compared to our recently published structure of the full-length S protein in the closed prefusion conformation^28^. The structure of the complex with three ACE2 bound is not symmetrical, as the distances between the C-termini of the three ACE2s (residue Tyr613) are 107Å, 109Å and 120Å, respectively (Fig. S4). This distance in the complex with two ACE2s bound is 110Å. These observations suggest that there is a modest degree of freedom for the up conformation of RBD when ACE2 is bound. All the substrate binding sites of the bound ACE2s face away from the threefold axis of the S trimer (Fig. S4), incompatible with the structure of the full length ACE2 dimer in complex with the amino acid transporter B^0^AT1, in which the two active sites of the two protomers are facing each other^18^. If the B^0^AT1-bound ACE2 dimer is indeed the form recognized by SARS-CoV-2, it appears that only one ACE2 protomer in the dimer can bind one RBD in an S trimer unless there are unexpectedly large structural rearrangements in either ACE2 or S.

**Figure 1.**
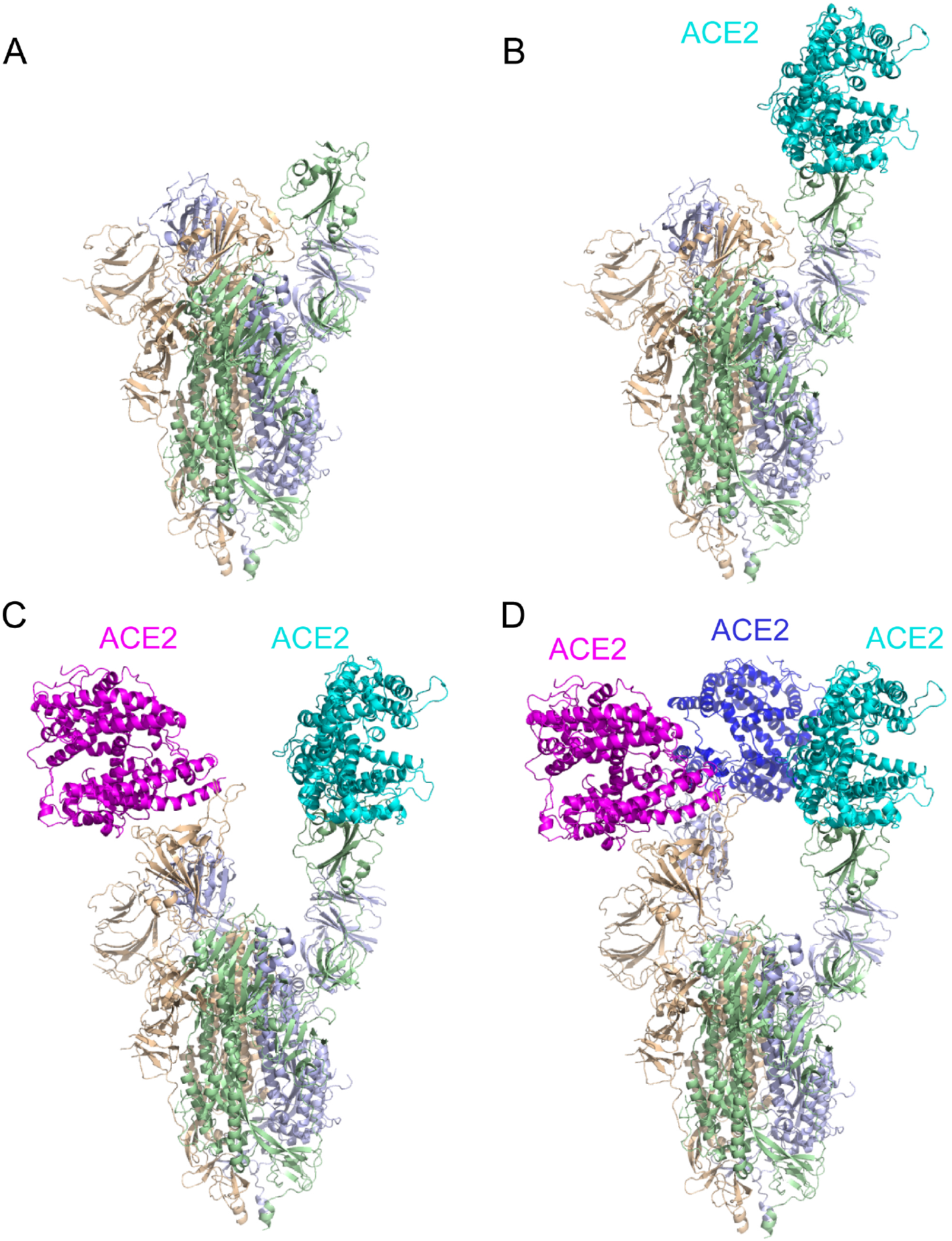
Cryo-EM structures of the ACE2-soluble S trimer complexes. Four distinct classes were identified and refined from a sample prepared by mixing a monomeric ACE2 and the stabilized soluble SARS-CoV-2 trimer. (A) The structure of the S trimer without ACE2 in a conformation with one RBD up was modeled based on a 3.6Å density map. Three protomers are colored in green, blue and wheat, respectively. (B)-(D) The structures of the S trimer in complex with one ACE2 (3.6Å), two ACE2 (3.7Å) and three ACE2 (3.4Å), respectively. ACE2 is colored in dark blue, cyan or magenta.

### Design of ACE2 variants to enhance its binding affinity to SARS-CoV-2 S trimer

Measurements of the binding kinetics of soluble monomeric ACE2 (ACE2_615_; Fig. 2) to the SARS-CoV-2 S trimer shows a relatively fast dissociation rate^28^, limiting its ability to compete with the membrane bound ACE2 on the surface of a target cell. We therefore sought to enhance the effective affinity by creating a trimeric form of soluble ACE2. We fused a trimerization foldon tag, derived from bacteriophage T4 fibritin^30^, to the C-terminal end of the ACE2 peptidase domain (residue 615) through a 11-residue flexible linker, a construct we refer to as ACE2_615_-foldon (Fig. 2). We have also created another version (ACE2_615_-LL-foldon) with a slightly longer linker (LL) with 13 residues between ACE2 and the foldon tag to assess its impact on binding. To further strengthen the interaction between ACE2 and the RBD, we introduced mutations guided by high-resolution structures (Fig. S5; ref^31^), at three different positions in the ACE2-RBD interface - T27, H34 and K353, respectively. Substitution with a bulky hydrophobic residue at each of these sites may enhance hydrophobic interactions between the two proteins and slow the dissociation (Fig. S5). We designed five mutants, T27Y, T27W, H34W, K353Y and K353W, in the ACE2_615_-foldon background. Finally, for comparison, we also include two versions of dimeric forms, ACE2m_615_-Fc and ACE2_740_-Fc, both fused to an Fc domain of an immunoglobulin G (Fig. 2A). ACE2m615-Fc contains H374N and H378N mutations at its peptidase active site and ACE2740-Fc includes the neck domain that mediates dimerization in the full length ACE2^18^.

**Figure 2.**
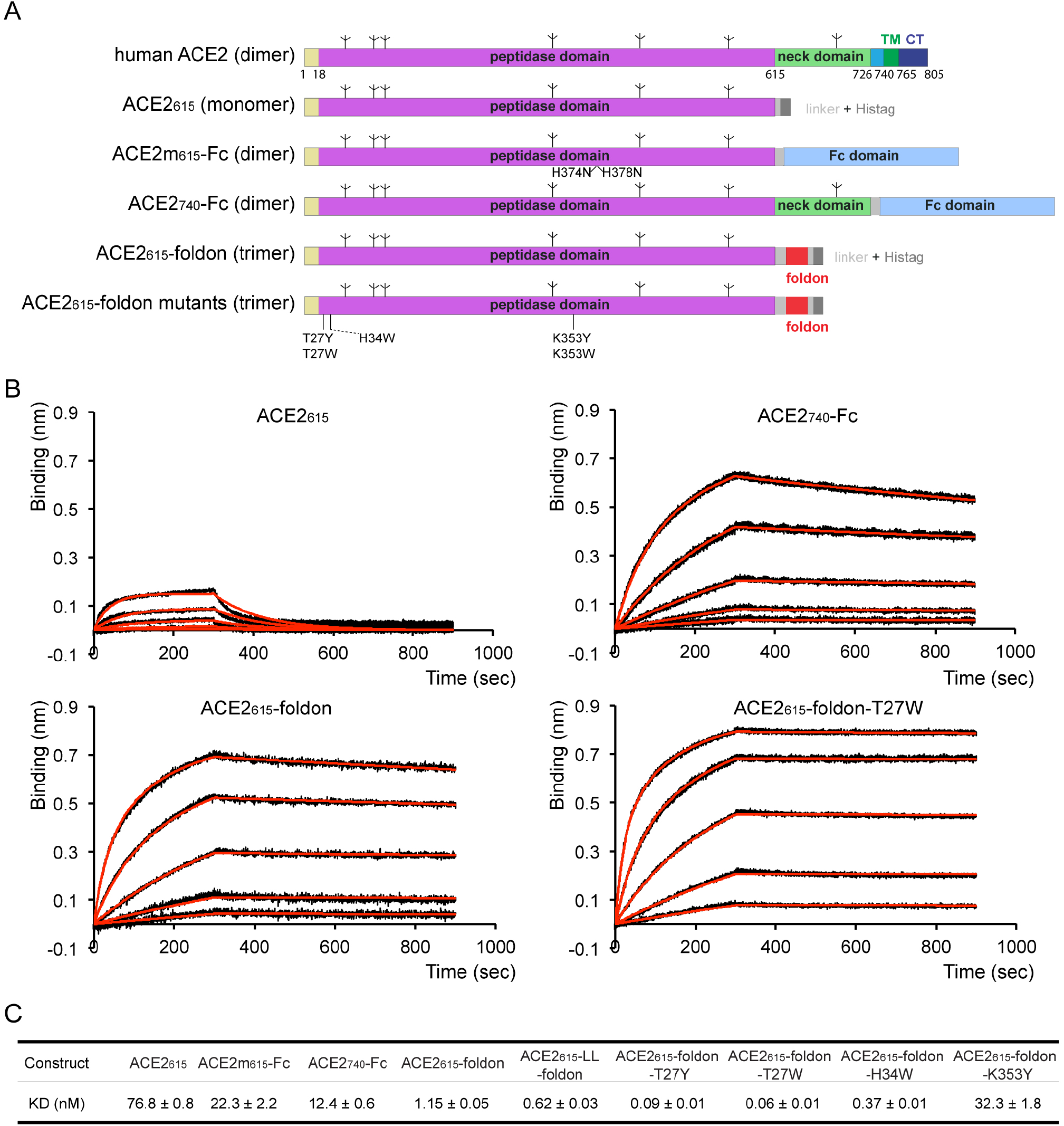
Design and characterization of ACE2 variants. (A) Schematic representation of the full-length human ACE2. Various segments include: catalytic peptidase domain, neck domain; TM, transmembrane anchor; CT, cytoplasmic tail; and tree-like symbols for glycans. Expression constructs of various forms of ACE2 used in this study: ACE2_615_, an inactive peptidase domain with mutations at the active site (H374N and H378N) fused with a C-terminal histag via a flexible linker; ACE2m_615_-Fc, the peptidase domain fused to a Fc fragment of an immunoglobulin G at the C-terminus; ACE2_740_-Fc, the peptidase and neck domains fused to a Fc fragment at the C-terminus; ACE2_615_-foldon, the peptidase domain fused to a trimerization tag-foldon, followed by a C-terminal histag; ACE2_615_-foldon mutants, single mutations (T27Y, T27W, H34W, K353Y and K353W) were introduced in the context of ACE2_615_-foldon construct. (B) Binding of ACE2 variants to the stabilized soluble S trimer by bio-layer interferometry (BLI). The S protein was immobilized to AR2G biosensors, which were dipped into the wells containing ACE2 at various concentrations (1.852-150 nM for ACE2_615_, 0.926-75 nM for ACE2_740_-Fc and 0.617-50 nM for all the ACE2_615_-foldon variants). Binding kinetics was evaluated using a 1:1 Langmuir binding model for the monomeric ACE2_615_ and a bivalent model for all other oligomeric ACE2. The sensorgrams are in black and the fits in red. All experiments were repeated at least twice with essentially identical results. (C) Summary of binding constants derived from the BLI experiments.

To produce the soluble recombinant ACE2 and its variants, we transfected HEK293 cells with the expression constructs of the monomeric and trimeric forms containing a C-terminal his tag and purified the proteins by Ni-NTA and gel filtration chromatography. The two dimeric forms were purified by protein G resin followed by gel filtration chromatography. While the monomeric and dimeric forms of soluble ACE2 were mostly secreted into cell medium, as judged by western blot, the trimeric ACE2_615_-foldon and its mutants were largely retained inside the cells. We therefore purified the secreted monomer and dimers from the cell supernatants and all the trimers from the cell lysates. Most proteins eluted from a size-exclusion column as a major symmetrical peak, regardless their secretion status (Fig. S6). The ACE2_740_-Fc protein containing the dimerizing neck domain appeared to aggregate substantially more than other constructs. The ACE2_615_-foldon K353W mutant aggregated completely, and we therefore did not pursue this construct any further. Only the fractions from the major peak for each construct were pooled and used for subsequent analyses. We also compared the secreted ACE2_615_-foldon with the form purified from the cells and their biochemical properties were almost identical (Fig. S7).

### Binding to SARS-CoV-2 soluble S trimer

We next measured binding of these recombinant ACE2 constructs to the stabilized soluble S trimer by bio-layer interferometry (BLI). As shown in Figs. 2B and S8, the monomeric ACE2_615_ had a fast dissociation rate and a K_D_ of 77 nM, consistent with the measurement that we reported recently using surface plasmon resonance (SPR; ref^28^). The dimeric ACE2m_615_-FC bound slightly more tightly (K_D_ ~22 nM). The dimeric ACE2_740_-Fc also bound more strongly than did the monomer (K_D_ ~12 nM), although the dimer formed by the neck domain is not compatible with two ACE2 peptidase domains interacting with two distinct RBDs in a single S trimer (Fig. S4). A possible explanation is that the neck domain mediated dimerization is not very strong and that the ACE2 peptidase domains are much more flexible than what the full-length ACE2 structure has indicated^18^, particularly in the absence of B^0^AT1. The trimeric ACE2_615_-foldon interacted with the S trimer much more strongly than any of the monomeric or dimeric forms, with a K_D_ of 1.2 nM. ACE2_615_-LL-foldon with a longer linker between ACE2 and foldon showed an additional modest affinity enhancement (K_D_ ~0.62 nM). The two interface mutants, ACE2_615_-foldon-T27W and ACE2_615_-foldon-T27Y, bound substantially more tightly than did the trimeric wild-type ACE2, with K_D_s of ~60 and ~90 pM, respectively. While the H34W afforded a slight affinity increase, the K353Y mutation decreased affinity by more than 25-fold. Overall, these data show that our structure-guided design to increase the affinity of ACE2 to the S trimer, by trimerizing the receptor and by modifying the interface, has indeed been effective.

### ACE2 peptidase activity and AT1R activation

We performed two independent assays to determine the enzymatic activity of these ACE2 constructs. First, we directly measured the peptidase activity using a synthetic peptide substrate that releases a free fluorophore upon ACE2 cleavage. In Fig. 3A, concentrations of all the proteins were normalized based on the number of active sites, and the fluorophore release was monitored continuously up to ~40 min. While all the trimeric forms showed essentially the same specific activity, the monomeric ACE2_615_ and the dimeric ACE2_740_-Fc had lower specific activities. The ACE2m_615_-Fc was inactive due to the mutations at the active site. Thus, all these ACE2 constructs with the wildtype sequence at the active site retained their wildtype peptidase activity.

**Figure 3.**
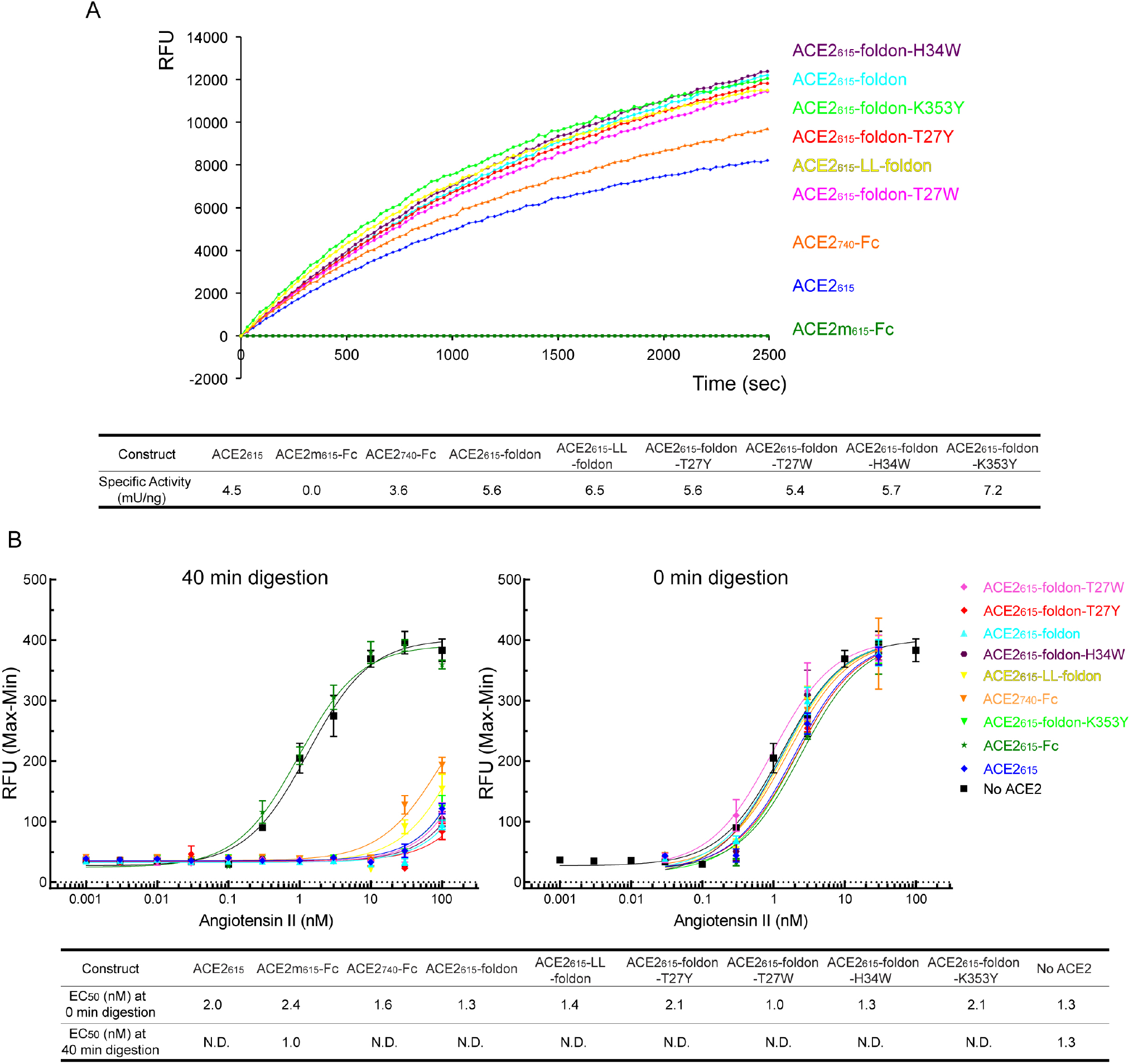
ACE2 peptidase activity and negative regulation of Ang II receptor type 1 activation. (A) Peptidase activity of the ACE2 variants were measured by detecting free fluorophore released from a synthetic peptide substrate. A time-course experiment was performed and specific activity was calculated. The experiment has been repeated twice with similar results. (B) Ang II peptide was treated with various ACE2 variants before adding to the cells expressing Ang II receptor type 1 (AT1R) at different concentrations. AT1R activation was quantified by changes in the intracellular calcium concentration. Samples quenched at time 0 were used as controls. The y-axis is Ratio (Max-Min) (Peak fluorescent intensity - baseline fluorescent intensity). The EC_50_ values were also summarized.

To further support this conclusion, we next tested the ability of the ACE2 constructs to block Ang II-induced activation of AT1R. In Fig. 3B, an Ang II peptide was first directly incubated with various ACE2 proteins and then added to HEK293 cells transfected with an AT1R expression construct. Activation of AT1R was monitored by changes in the intracellular calcium concentration. When the digestion reaction was quenched by EDTA at time 0 as a control, all the mixtures with different ACE2 proteins could efficiently activate AT1R, suggesting that nothing in our protein preparations inhibited Ang II-mediated AT1R activation. In contrast, when the digestion was allowed to proceed for 40 min, all AEC2 constructs except for the inactive ACE2m_615_-Fc effectively blocked AT1R activation, presumably by converting Ang II to Ang 1–7, in agreement with the peptidase activity results.

### Inhibition of SARS-CoV-2 infectivity in cell culture

We used three different assays to assess the neutralization potency of the ACE2 constructs in blocking SARS-CoV-2 infection. The circulating strain during the early days of the pandemic contained a D614 residue in its S protein, but it has subsequently been replaced by an emerging strain harboring a G614 substitution^32^. It has been difficult to generate pseudotyped viruses with the full-length S from the D614 strain. We first used an MLV-based pseudovirus assay with a D614 S construct lacking 19 residues of the cytoplasmic tail, which incorporates efficiently into pseudoviruses. In Fig. 4A, the monomeric ACE2_615_ showed the lowest potency with an IC_50_ value of 24.1 μg/ml. The two dimeric forms, ACE2m_615_-Fc and ACE2_740_-Fc, and the trimeric mutant ACE2_615_-foldon-K353Y had greater potency, with IC_50_ values ranging from 1.2 to 6.3 μg/ml. The two trimeric forms, ACE2_615_-foldon and ACE2_615_-LL-foldon, and the trimeric mutant ACE2_615_-foldon-H34W neutralized with even greater potency and an IC_50_ value around 0.6 μg/ml. The most potent inhibitors were ACE2_615_-foldon-T27W and ACE2_615_-foldon-T27Y, which had IC_50_ values of 0.21 and 0.25 μg/ml, respectively. Thus, the neutralization potency of these ACE2 constructs correlates strictly with their binding affinity, suggesting that the interaction between ACE2 and S is the principal determinant of neutralization of the virus pseudotyped with the CT-truncated S (D614). Neutralization by these ACE2 proteins in the HIV-based pseudovirus assay using a full-length S derived from the G614 circulating strain showed a very similar pattern with ACE2_615_ the weakest and ACE2_615_-foldon-T27W and ACE2_615_-foldon-T27Y the most potent (Fig. 4B). Furthermore, when they were analyzed by a plaque assay with an authentic SARS-CoV-2, the neutralization pattern was almost identical to that from the MLV-based assay (Fig. 4C). The IC_50_ values for ACE2_615_-foldon-T27W and ACE2_615_-foldon-T27Y were 0.08 and 0.14 μg/ml, respectively. These results indicate that the engineered ACE2 constructs are very potent agents for blocking SARS-CoV-2 infection in cell culture.

**Figure 4.**
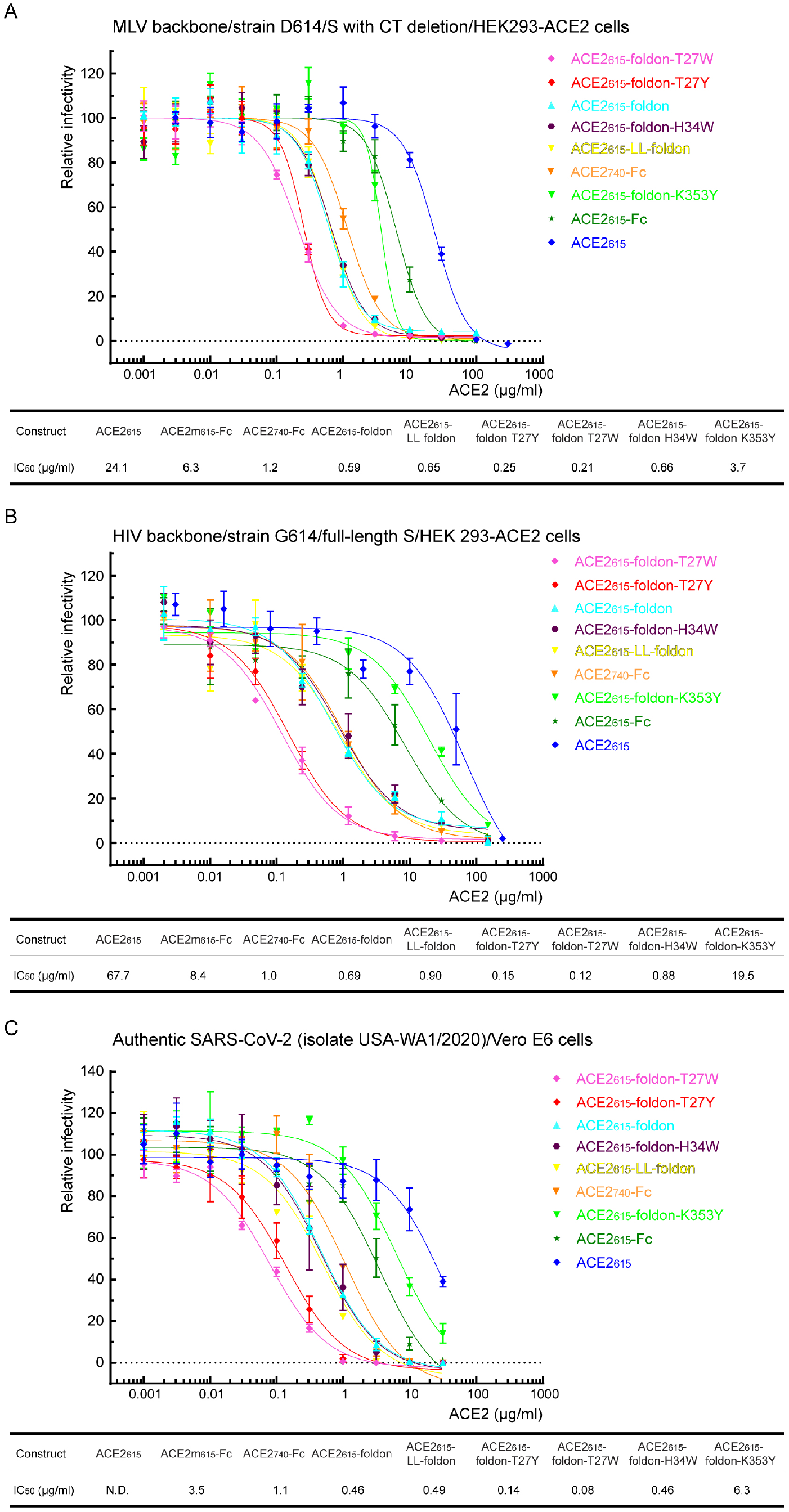
Inhibition of SARS-CoV-2 pseudoviruses and authentic viruses by ACE2 variants. (A) Serial dilutions of each ACE2 variant were tested for inhibition against an MLV-based pseudotyped virus using a SARS-CoV-2 S construct containing D614 and a CT deletion in a single-round infection of HEK293-ACE2 cells. IC_50_ values were derived from curve fitting. The experiments were repeated three times with similar results. (B) ACE2 variants were tested for inhibition against an HIV-based pseudotyped virus using a full-length SARS-CoV-2 S construct containing G614 in a single-round infection of HEK293-ACE2 cells. IC_50_ values were derived from curve fitting. The experiments were repeated twice with similar results. (C) Serial dilutions of each ACE2 variant were tested for inhibition against an authentic SARS-CoV-2 S virus (isolate USA-WA1/2020) infecting Vero E6 cells. IC_50_ values were derived from curve fitting. The experiments were repeated three times with similar results.

## Discussion

A recombinant human ACE2, named APN01, is currently under evaluation as a treatment for COVID-19 in a phase 2 clinical trial (NCT04335136), primarily based on the favorable results from a previous phase 1 safety and tolerability trial (NCT00886353) in a small number of healthy individuals^33^, as well as on the recent evidence that the protein blocks SARS-CoV-2 infection effectively *in vitro*^26^. APN01 is a soluble ACE2 construct expressing residues 1-740 and probably dimerizes by the neck domain^34^, like ACE2_740_-Fc used in our study. We demonstrate here that our best trimeric ACE2 variant, ACE2_615_-foldon-T27W, has >200-fold higher binding affinity for the soluble SARS-CoV-2 S trimer, and ~5-fold and ~13-fold higher neutralization potency against pseudoviruses and authentic viruses, respectively, than does ACE2_740_-Fc, while its peptidase activity and ability to block AT1R activation remain essentially unchanged. Using a deep mutagenesis screening approach, a recent study has identified a dimeric ACE2 variant containing multiple mutations, which led to higher affinity binding to the RBD, but also a substantial loss in the catalytic activity (4-8 fold decrease) than the parental construct with the wildtype sequence^35^. One of the mutations from the mutagenesis screening is T27Y, coinciding with our structure-based design. Our approach also distinguishes the S protein binding and the peptidase activity, which can be manipulated separately to maximize the therapeutic benefits of an ACE2 construct.

Although the molecular mechanism by which a soluble ACE2 blocks SARS-CoV-2 infection as a decoy receptor is obvious, its protective role against lung injury – a hallmark of severe COVID-19 cases – appears to be more complicated in humans than in animal models. ACE2 knockout mice have more severe ARDS symptoms than do wildtype mice, while ACE2 overexpression appears to be protective^23^. Moreover, administration of recombinant ACE2 reduces severity of lung injury in mice caused by respiratory syncytial virus or influenza virus^24,25^. In humans, rhACE2 was well tolerated with a short half-life^33^, but its infusion did not appear to ameliorate ARDS at least in a small number of patients^36^. The ongoing phase 2 clinical trial with expected 200 participants may provide much needed information regarding the therapeutic potential of recombinant ACE2 in the context of COVID-19. Structure-based modifications presented here may help augment therapeutic efficacy while suppressing adverse effects, if any.

The safety in humans of the foldon trimerization tag, derived from the bacteriophage T4 fibritin^30^, has been demonstrated by vaccine trials against HIV-1 and SARS-CoV-2 in clinical settings^37,38^. However, a dose in mg/kg body weight of the trimeric ACE2 proteins as a therapeutics is likely much greater than that used as a vaccine (for example, 50-250 μg of foldon stabilized HIV-1 gp140 protein/injection; ref^37^). If the foldon tag induces unacceptable levels of side effects at a high dose in animals or humans, other trimerization domains, such as those in abundant human collagens^39^, can be considered. Further improvements of these ACE2-based therapeutic candidates include modifications to enhance protein stability by introducing additional disulfide bonds (which may reduce the catalytic activity), to modulate peptidase activity by mutating residues in or near the active site, and to increase its in vivo residence time in circulation, by strategies such as PEGylation^40^.

The structure of the membrane-bound ACE2 dimer formed by the neck domain is not compatible with a binding mode of two protomers interacting with two RBDs from a single S trimer, as depicted in Fig. S4. Much stronger binding of ACE2_740_-Fc to the S trimer, as well as greater neutralization potency than those of monomeric ACE2_615_ and even dimeric ACE2m_615_-Fc clearly indicate avidity, suggesting that the ectodomain of either ACE2 or SARS-CoV-2 S protein has much greater flexibility than the cryo-EM structures imply ^18,28,41,42^. If the multivalency of the trimeric S protein on the surface of virion and the dimeric ACE2 on the host cells indeed plays an important role during viral attachment, the binding affinity for the virus to latch on the target cells would be much stronger than the values measured using monomeric ACE2^41^. It may help explain the unexpected transmission efficiency of SARS-CoV-2 leading to a pandemic on a surprising scale and raise the hurdle for antivirals to effectively compete with ACE2 for S binding. Trimeric ACE2 variants exerting even greater avidity than the dimeric form on the host cells may have a competitive edge over other RBD-targeting inhibitors, such as monoclonal antibodies, with similar binding affinity.

## Acknowledgments

We thank Gary Frey for generous advice, Sarah Sterling, Richard Walsh Jr. and Shaun Rawson for technical support, and Stephen Harrison for critical reading of the manuscript. EM data were collected at the Harvard Cryo-EM Center for Structural Biology of Harvard Medical School. This work was supported by NIH grants AI147884 (to B.C.), AI147884-01A1S1 (to B.C), AI141002 (to B.C.), AI127193 (to B.C. and James Chou), a COVID19 Award by Massachusetts Consortium on Pathogen Readiness (MassCPR; to B.C.), as well as a Fast grant by Emergent Ventures (to B.C.).

## Author Contribution

B.C. and T.X. conceived the project. T.X. designed, expressed and purified ACE2 variants with help from H.P. and Y.C.. T.X. also performed binding experiments and enzymatic assays. J.L. carried out the neutralization assays using the MLV-based pseudoviruses, also designed and performed the AT1R activation experiments with contributions from S.L.. J.Z. determined the cryo-EM structures of the ACE2-S complexes. R.I.J., L.G.A.M., N.S. and A.G. carried out the neutralization assays using SARS-CoV-2. C.L.L. and M.S.S performed the neutralization assays using the HIV-based pseudoviruses. Y.C. also designed and produced the soluble S trimer. S.R.V. contributed to cell culture for protein production. B.D.Q. and M.F. created the expression constructs for the dimeric ACE2 variants. All authors analyzed the data. B.C. and T.X. wrote the manuscript with input from all other authors.

## Methods

### Protein expression and purification

A synthetic gene encoding an human ACE2 fragment (residues 1-615) fused with a C-terminal 6xHis tag was generated by GenScript (Piscataway, NJ) and cloned into pCMV-IRES-puro expression vector (Codex BioSolutions, Inc, Gaithersburg, MD) to create the construct pACE2_615_. To construct a trimeric ACE2 variant, a DNA fragment encoding a foldon trimerization tag was inserted between the ACE2 fragment and the His tag by restriction digestion and DNA ligation to give the plasmid pACE2_615_-foldon. Site-specific mutations were introduced to the ACE2_615_-foldon construct by PCR following standard protocols of site-directed mutagenesis. All the ACE2 variants were expressed in HEK 293F cells by transient transfection using Opti-MEM (Gibco-Thermo Fisher Scientific, Waltham, MA). After incubation for 4 days at 37°C with 5.5% CO_2_, the transfected cells were harvested by centrifugation at 2,524 xg for 30 minutes.

For the monomeric ACE2_615_ protein, the cell supernatant was collected by centrifugation and loaded onto a column packed with Ni-NTA agarose beads (Qiagen, Hilden, Germany). The column was washed with a buffer containing 20 mM Tris-HCl, pH 7.5, and 300 mM NaCl. The protein was eluted using a buffer containing 100 mM imidazole, and further purified by gel filtration chromatography on a Superdex 200 Increase 10/300 GL column (GE Healthcare, Chicago, IL)

To purify the dimeric ACE2_615_-Fc and ACE2_740_-Fc proteins, the cell supernatant was collected and loaded to a column packed with GammaBind Plus Sepharose beads (GE Healthcare). The column was washed with PBS. The protein was eluted using 100 mM glycine (pH 2.5) and neutralized immediately with 2 M Tris-HCl (pH 8.0). The eluted protein was further purified by gel filtration chromatography on a Superdex 200 Increase 10/300 GL column.

For all the ACE2_615_-foldon variants, which were not secreted efficiently, the cell pellet was resuspended in the lysis buffer (20 mM Tris-HCl, pH 7.5, 300 mM NaCl, 1% NP40, 20 mM imidazole) and rocked gently for 1 hour at 4°C, followed by spinning at 17,554 xg for 30 minutes to remove cell debris. The supernatant was loaded to a column packed with Ni-NTA agarose beads (Qiagen). The column was washed with a buffer containing 20 mM Tris-HCl, pH 7.5, 300 mM NaCl and 50 mM imidazole and the protein was eluted using a buffer containing 20 mM Tris-HCl, pH 7.5, 300 mM NaCl and 300 mM imidazole. The eluted protein was further purified by gel filtration chromatography on a Superdex 200 Increase 10/300 GL column.

To produce a stabilized ectodomain of SARS-CoV-2 S trimer protein, a synthetic gene (kindly provided by Dr. Dan Barouch), encoding residues 1-1208 with the furin cleavage site (residues 682–685) replaced by a “GGSG” sequence, residues K986 and V987 substituted by prolines, and addition of a foldon trimerization tag followed by a C-terminal 6xHisTag, was cloned into the vector pCMV-IRES-puro. The expression construct was transiently transfected in HEK 293T cells using polyethylenimine (Polysciences, Inc, Warrington, PA). Protein was purified from cell supernatants using Ni-NTA resin (Qiagen), the eluted fractions containing S protein were pooled, concentrated, and further purified by gel filtration chromatography on a Superose 6 column (GE Healthcare).

### Cryo-EM sample preparation and data collection

To prepare cryo grids, 3.5 μl of the freshly prepared mixture of the soluble S trimer and monomeric ACE2 (1:3 molar ratio) at ~1 mg/ml was applied to a 1.2/1.3 Quantifoil grid (Quantifoil Micro Tools GmbH, Germany), which had been glow discharged with a PELCO easiGlow™ Glow Discharge Cleaning system (Ted Pella, Inc., Redding, CA) for 60 s at 15 mA. Grids were immediately plunge-frozen in liquid ethane using a Vitrobot Mark IV (Thermo Fisher Scientific), and excess protein was blotted away using grade 595 filter paper (Ted Pella, Inc.) with a blotting time of 4 s, a blotting force of −12 at 4°C in 100% humidity. The grids were first screened for ice thickness and particle distribution using a Talos Arctica transmission electron microscope (Thermo Fisher Scientific), operated at 200 keV and equipped with a K3 direct electron detector (Gatan), at the Harvard Cryo-EM Center for Structural Biology. For data collection, images were acquired with selected grids using a Titan Krios transmission electron microscope (Thermo Fisher Scientific) operated at 300 keV with a BioQuantum GIF/K3 direct electron detector. Automated data collection was carried out using SerialEM version65 at a nominal magnification of 105,000× and the K3 detector in counting mode (calibrated pixel size, 0.825 Å) at a exposure rate of ~14.8 electrons per physical pixel per second. Each movie had a total accumulated electron exposure of 50 e/Å2 fractionated in 50 frames of 50 ms. Datasets were acquired using a defocus range of 1.5-2.6 μm.

### Image processing, 3D reconstructions and model building

Drift correction for cryo-EM images was performed using MotionCor2^43^, and contrast transfer function (CTF) was estimated by CTFFIND4^44^ using motion-corrected sums without dose-weighting. Motion corrected sums with dose-weighting were used for all image processing. CrYOLO^45^ was used for particle picking, and RELION3.0.8^46^ was used for 2D classification, 3D classification and refinement. A total of 407,761 particles were extracted from 4,292 images. The selected particles were subjected to 2D classification, giving a total of 261,799 good particles. A low-resolution negative-stain reconstruction of the sample was low-pass-filtered to 40Å and used as an initial model for 3D classification in C1 symmetry. One class containing 32,685 particles appeared to represent the free S trimer with no ACE bound was further refined in C1 symmetry, giving a reconstruction at 3.6Å resolution. Another major class with ~49% of the selected particles showing density for ACE2 was refined in C1 symmetry and subsequently subjected to CTF refinement, Bayesian polishing and particle subtraction by masking out the ACE2-RBD density, followed by 3D classification without alignment in six classes. Whole particles were re-extracted based on the six classes from the masked local classification and refined further, revealing different stoichiometry for ACE2 binding (one ACE2 per S trimer, two ACE2 per S trimer, and three ACE2 per S trimer). Three best maps representing each type of complexes were chosen and further refined in C1 symmetry after CTF refinement and Bayesian polishing, leading to one reconstruction of the complex with one ACE2 bound at 3.6Å resolution from 15,964 particles; another reconstruction of the complex with two ACE2 bound at 3.7Å resolution from 13,515 particles and a third reconstruction of the complex with three ACE2 bound at 3.4Å resolution from 26,298 particles. Reported resolutions are based on the gold-standard Fourier shell correlation (FSC) using the 0.143 criterion. All density maps were corrected from the modulation transfer function of the K3 detector and then sharpened by applying a temperature factor that was estimated using post-processing in RELION. Local resolution was determined using RELION with half-reconstructions as input maps.

The initial templates for model building used the stabilized SARS-CoV-2 S ectodomain trimer structure (PDB ID: 6vyb) and ACE2 from the ACE2-B0AT1 complex structure (PDB ID: 6M17). Several rounds of manual building were performed in Coot. Iteratively, refinement was performed in both Phenix^47^ (real space refinement) and ISOLDE ^48^, and the Phenix refinement strategy included rigid body fit, minimization_global, local_grid_search, and adp, with rotamer, Ramachandran, and reference-model restraints, using 6vyb and 6M17 as the reference model. The refinement statistics are summarized in Table S1.

### Binding assay by bio-layer interferometry (BLI)

Binding of ACE2 variants to the soluble S trimer was measured using an Octet RED384 system (ForteBio, Fremont, CA). Each ACE2 protein was diluted using the running buffer (PBS, 0.005% Tween 20, 0.25 mg/ml BSA) and transferred to a 96-well plate. The soluble S protein was immobilized to Amine Reactive 2^nd^ Generation (AR2G) biosensors (Fortebio), following a protocol recommended by the manufacturer. After equilibrating in the running buffer for 5 minutes, the sensors with immobilized S protein were dipped in the wells containing the ACE2 protein at various concentrations (1.852-150 nM for ACE2_615_; 0.926-75 nM for ACE2_615_-Fc and ACE2_740_-Fc; 0.617-50 nM for all the ACE2_615_-foldon variants) for 5 minutes to measure the association rate. The sensors were then dipped in the running buffer for 10 minutes to determine the dissociation rate. Control sensors with no S protein were also dipped in the ACE2 solutions and the running buffer as references. Recorded sensorgrams with background subtracted from the references were analyzed using the software Octet Data Analysis HT Version 11.1 (Fortebio). The curves for monomeric ACE2 were fit to a 1:1 binding model, while those for the oligomeric ACE2 variants were fit to a bivalent binding model.

### ACE2 peptidase activity assay

The catalytic activity of the ACE2 variants was measured by detecting a free fluorophore 7-methoxycoumarin-4-acetic acid (MCA) released from a synthetic peptide substrate, using an ACE2 activity kit (BioVision, Milpitas, CA). The ACE2_615_ and ACE2_615_-foldon variants were diluted to 0.25 μg/ml using the assay buffer from the kit. The ACE2_615_-Fc and ACE2_740_-Fc proteins were diluted to 0.38 and 0.30 μg/ml, respectively, to keep the same number of the active sites as other ACE2 variants. 50 μl of diluted protein was set in the 96-well plate. Immediately before recording fluorescence signals, 50 μl substrate diluted in the assay buffer, following a protocol recommended by the manufacturer, was added to each well. Fluorescence signals were recorded in a kinetic mode by a Flexstation 3 Multi Mode Microplate Reader (Molecular Devices, San Jose, CA). The specific activity was calculated as the amount of the released fluorophore divided by the reaction time and the amount of the ACE2 protein using the data within the initial linear phase, as described in the protocol provided by the manufacturer. To determine the initial linear phase, fluorescence signals were recorded with 1.25, 0.25 and 0.125 μg of ACE2_615_ protein, respectively, reaching maximum after the substrates were completely cleaved. Data from the first 2 minutes within the linear phase with signals less than 10% of the maximum were used for the calculation. The amount of released MCA was derived from the increase of the fluorescence signal divided by the slope of the MCA standard curve.

### Inhibition of Ang II-induced AT1R activation

To treat the Ang II peptide with each ACE2 variant, 2 μl of ACE2 protein at 0.5 mg/ml were added to 198 μl of an assay buffer (1xPBS, 40 mM Tris-HCl, pH6.8, 20 μM ZnCl_2_) containing 65 μM Ang II peptide. The reactions were incubated at 37°C for 40 min, and then quenched by addition of 50 μl of 0.5 M EDTA. The final concentration of Ang II peptide was 52 μM. As a time 0 control, 198 μl of the assay buffer containing 65 μM Ang II was incubated with EDTA at 37°C first, followed by addition of 2 μl of each ACE2 protein (0.5 mg/ml).

Changes in the intracellular calcium concentration in AT1R-expressing cells when induced by Ang II peptide were measured to monitor the activation of the receptor. Briefly, HEK293 cells were transfected with pCMV-AT1R-IRES-Puro gene using Lipofectamine 3000 reagent (Thermo Fisher Scientifics). Approximately 24 hours posttransfection, the cells were transferred into a 384-well black clear plate at a density of 1.2×10^4^ cells/well in 20 μl culture medium. On day 4, 20 μl of 1x Non-Wash Calcium Dye solution (CB-80500-301, Codex BioSolutions Inc) was added into each well. The cell plate was incubated at 37°C in a CO_2_ incubator for 1 hour. The pretreated ligands (Ang II peptide) at various concentrations (0.005-500 nM) were prepared in 1X HBSS with 20 mM HEPES (pH7.46). Fluorescent intensity in each well was recorded on an FDSS 7000 (Hamamatsu Corporation, Bridgewater, NJ) at the rate of 1 image/sec (Ex 480 nM and Em 540 nM) and the base line of each well was also recorded for 10 seconds. After the online addition of 10 μl of the prepared ligands (the final concentration of 0.001-100 nM), the fluorescent intensity of each well was recorded at the rate of 1 image/sec for additional 170 seconds.

### MLV-based pseudovirus assay

Murine Leukemia Virus (MLV) particles (all plasmids of the MLV components were kindly provided by Dr. Gary Whittaker at Cornell University and Drs. Catherine Chen and Wei Zheng at National Center for Advancing Translational Sciences, National Institutes of Health), pseudotyped with a SARS-CoV-2 S protein construct, were generated in HEK 293T cells, following a protocol described previously for SARS-CoV^49,50^. To enhance incorporation, C-terminal 19 residues in the cytoplasmic tail of the SARS-CoV-2 S protein containing D614 were deleted. To prepare for infection, 7.5×10^3^ of Expi-293F cells, stably transfected with a full-length human ACE2 expression construct, in 15 μl culture medium were plated into a 384-well white-clear plate coated with poly-D-Lysine to enhance cell attachment. On day 2, 12.5 μl of SARS-CoV-2 MLV pseudoviruses were mixed with 5 μl of each ACE2 variant at different concentrations (0.001-300 μg/ml) and incubated at 37°C for 1 hr. After the medium in each well containing the cells was removed, 17.5 μl of each ACE2-virus mixture were added. The plate was centrifuged at 54 xg for 15 min at 4°C and additional 7.5 μl of culture medium were then added. The total final volume in each well was 25 μl. The cells were then incubated at 37°C for 42 hr. Luciferase activities were measured with Firefly Luciferase Assay Kit (CB-80552-010, Codex BioSolutions Inc). IC_50_ values were calculated based on curve fitting in GraphPad Prism.

### HIV-based pseudovirus assay

Neutralization of HIV-based pseudovirus containing a full-length SARS-CoV-2 S protein was measured using a single-round infection assay in HEK 293T/ACE2 target cells. Pseudotyped virus particles were produced in 293T/17 cells (ATCC) by co-transfection of a plasmid encoding codon-optimized SARS-CoV-2 full-length S containing G614, a packaging plasmid pCMV ΔR8.2 expressing HIV gag and pol, and a luciferase reporter plasmid pHR’ CMV-Luc. All plasmids were kindly provided by Dr. Barney Graham (NIH, Vaccine Research Center). The 293T cell line stably overexpressing the human ACE2 protein was created by the Farzan group at Scripps Research Institute. For neutralization assays, serial dilutions of the ACE2 constructs were performed in duplicate followed by addition of pseudoviruses. Plates were incubated for 1 hour at 37°C followed by addition of 293T/ACE2 target cells (1×10^4^/well). Wells containing cells and pseudoviruses without ACE2 proteins or cells alone were positive and negative infection controls, respectively. Assays were harvested on day 3 using BrightGlo luciferase reagent (Promega, Madison, WI) and luminescence detected with a Victor luminometer (PerkinElmer, Waltham, MA). IC_50_ values are reported as the ACE2 protein concentration that inhibited 50% virus infection. All neutralization experiments were repeated twice with similar results.

### Neutralization of authentic SARS-CoV-2

ACE2 variants were serially diluted in Dulbecco’s Phosphate Buffered Saline (DPBS)(Gibco™) using half-log dilutions starting at 31,579 ng/ml. Dilutions were prepared in triplicate for each protein. Each dilution was incubated at 37°C in 5% CO_2_ for 1 hour with 1,000 plaque forming units/ml (PFU/ml) of SARS-CoV-2 (isolate USA - WA1/2020). Controls included Dulbecco’s Modified Eagle Medium (DMEM) (Gibco-Thermo Fisher Scientific) containing 2% fetal bovine serum (Gibco-Thermo Fisher Scientific) and antibiotic-antimycotic (Gibco-Thermo Fisher Scientific) only as a negative control, 1000 PFU/ml SARS-CoV-2 (USA-WA1/2020) incubated with DPBS, and 1000 PFU/ml SARS-CoV-2 incubated with DMEM. 200 μl of each dilution or control were added to confluent monolayers of NR-596 Vero E6 cells in triplicate and incubated for 1 hour at 37°C and 5% CO_2_. The plates were gently rocked every 5-10 minutes to prevent monolayer drying. The monolayers were then overlaid with a 1:1 mixture of 2.5% Avicel^®^ RC-591 microcrystalline cellulose and carboxymethylcellulose sodium (DuPont Nutrition & Biosciences, Wilmington, DE) and 2X Modified Eagle Medium (Temin’s modification, Gibco-Thermo Fisher Scientific) supplemented with 2X antibiotic - antimycotic, 2X GlutaMAX (Gibco-Thermo Fisher Scientific) and 10% fetal bovine serum. Plates were incubated at 37°C and 5% CO_2_ for 2 days. The monolayers were fixed with 10% neutral buffered formalin and stained with 0.2% aqueous Gentian Violet (RICCA Chemicals, Arlington, TX) in 10% neutral buffered formalin for 30 min, followed by rinsing and plaque counting. The half maximal inhibitory concentrations (IC_50_) were calculated using GraphPad Prism 8.

