## Supplemental Figures and Table for "A trimeric human angiotensin-converting enzyme 2 as an anti-SARS-CoV-2 agent in vitro"

### Supplementary materials

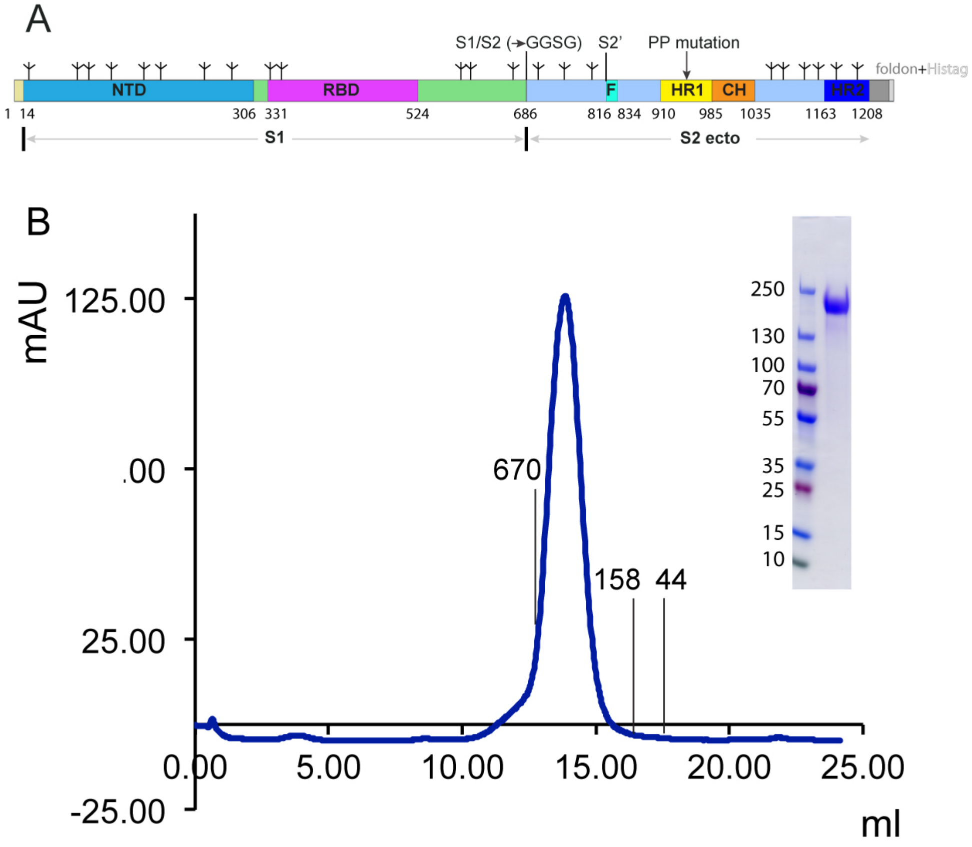


**Figure S1. Preparation of a stabilized soluble SARS-CoV-2 S trimer.** (A) Schematic representation of the expression construct of the soluble SARS-CoV-2 S protein. Segments of S1 and S2 ectodomain (S2ecto) include: NTD, N-terminal domain; RBD, receptor-binding domain; S1/S2, S1/S2 cleavage site; S2', S2' cleavage site; FP, fusion peptide; HR1, heptad repeat 1; CH, central helix region; HR2, heptad repeat 2; and tree-like symbols for glycans. The S1/S2 cleavage site (RRAR) was mutated to GGSG. Two mutations K986P and V987P were introduced and a trimerization tag –foldon fused to the C-terminal end to stabilize the prefusion conformation[27]. A C-terminal histag was included for protein purification. (B) The purified S protein was resolved by gel-filtration chromatography on a Superose 6 column and the pooled peak fractions were analyzed by Coomassie stained SDS-PAGE. The molecular weight standards include thyoglobulin (670 kDa), $\gamma$ -globulin (158 kDa) and ovalbumin (44 kDa).

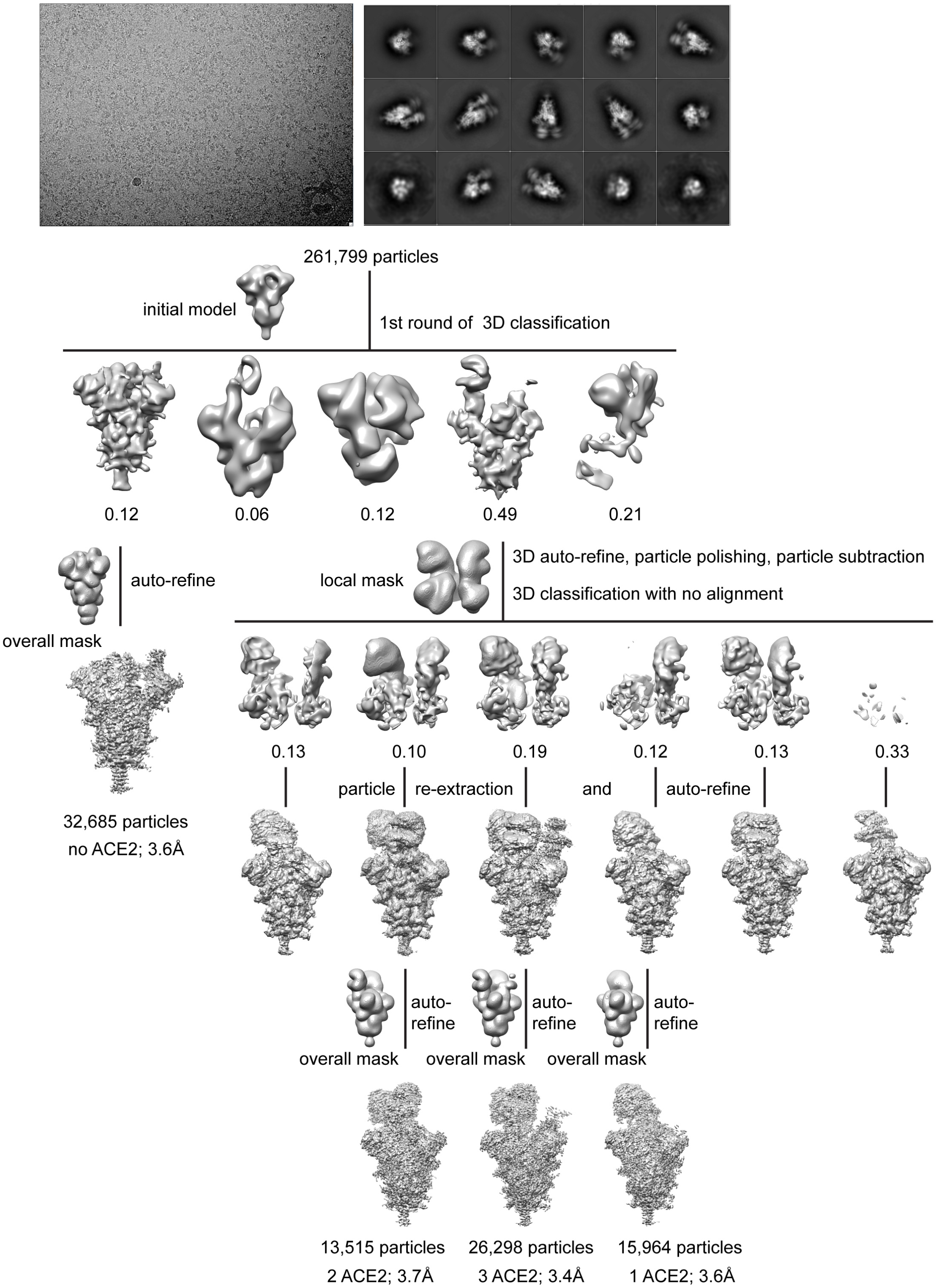


**Figure S2. Cryo-EM analysis of the ACE2-S complexes.** Top, representative micrograph, and 2D averages of the cryo-EM particle images showing secondary structural features. Bottom, data processing workflow for structures of the free S trimer (no ACE2), S trimer with one ACE2 bound (1 ACE2), S trimer with two ACE2 bound (2 ACE2), S trimer with three ACE2 bound (3 ACE2), as indicated.

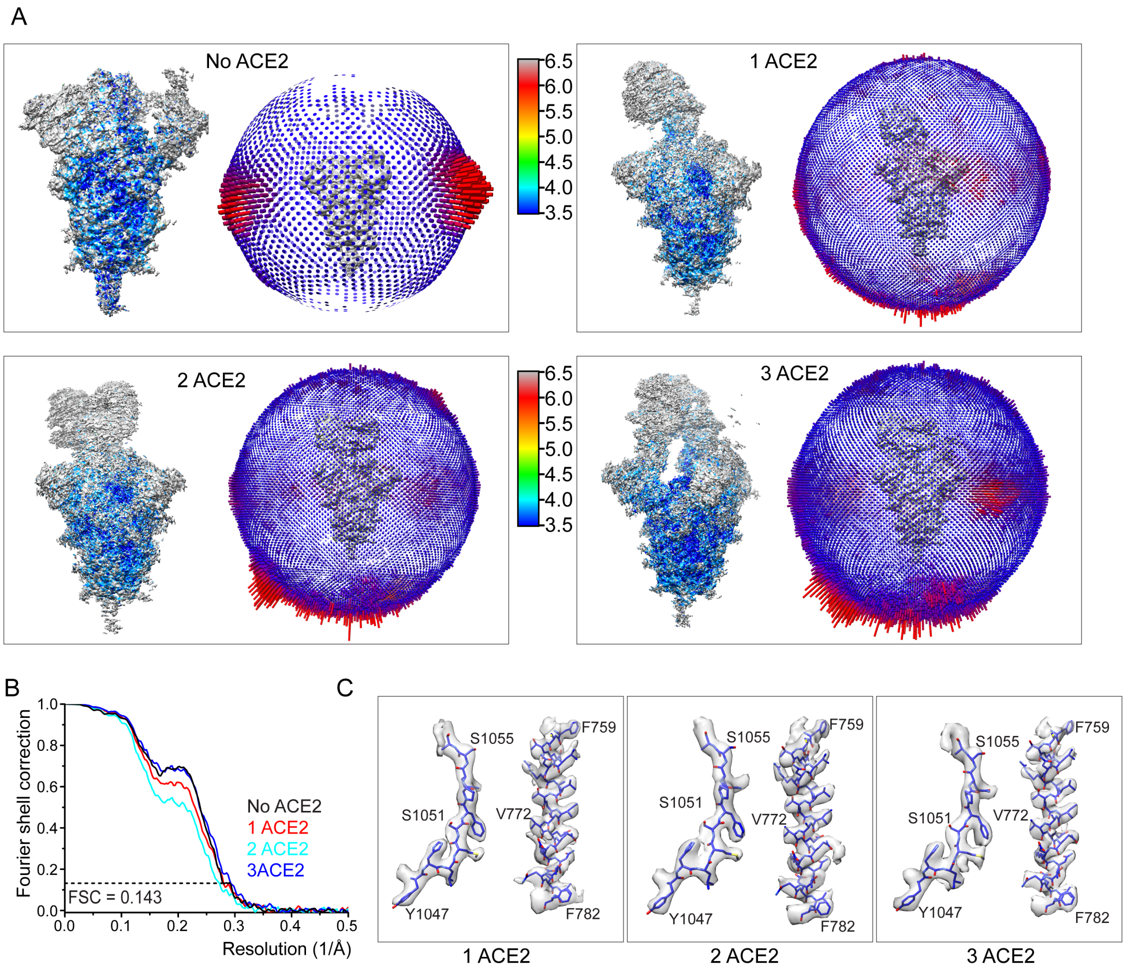


**Figure S3. Analysis of the 3D reconstructions of the S-ACE2 complexes.** (A) 3D reconstructions of the S trimer and its ACE2 complexes are colored according to local resolution estimated by RELION. Angular distribution of the cryo-EM particles used in the reconstruction is shown in the side view of the EM map. (B) Gold standard FSC curves of the refined 3D reconstructions. (C) Representative density in gray surface from the EM maps. No ACE2), the free S trimer; 1 ACE2, S trimer with one ACE2 bound; 2 ACE2, S trimer with two ACE2 bound; 3 ACE2, S trimer with three ACE2 bound.

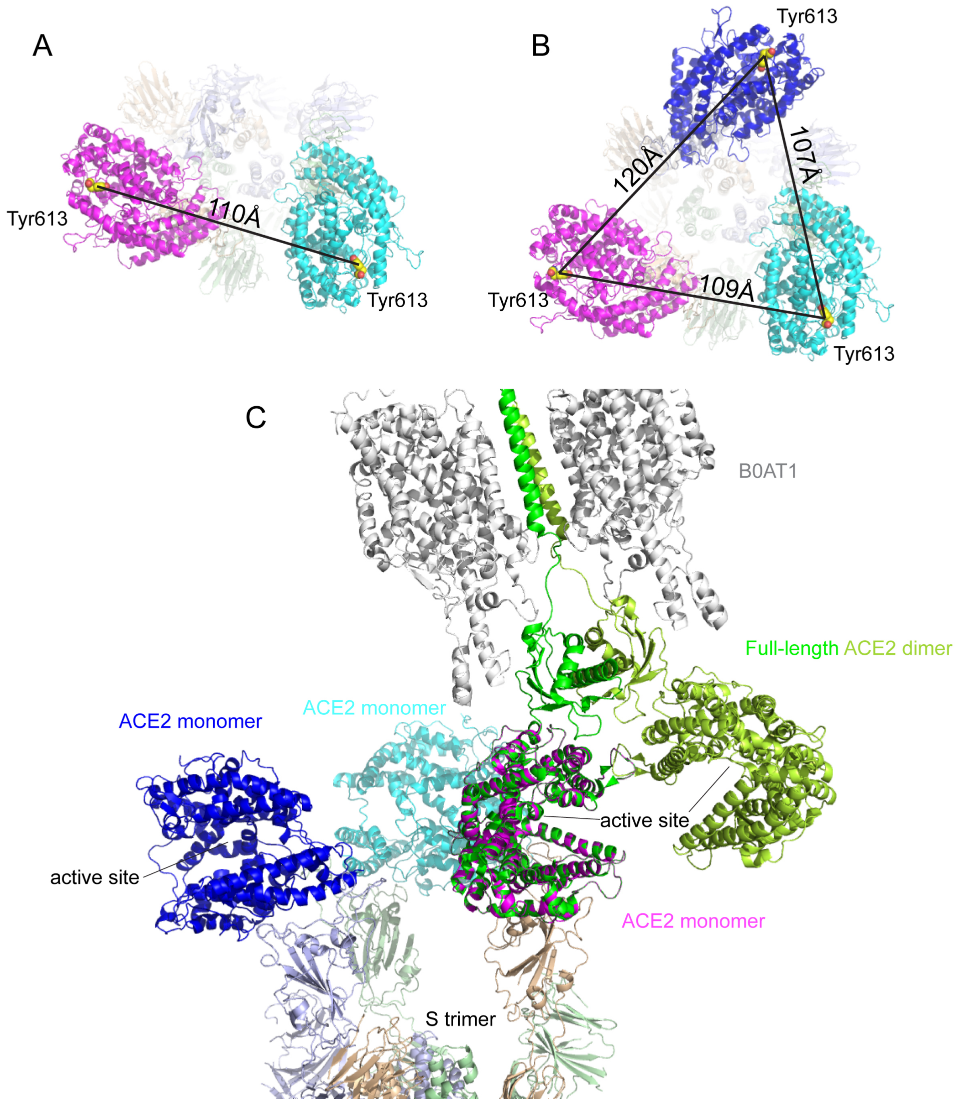


**Figure S4. ACE2-S interactions.** (A) The top view of the S trimer in complex with two monomeric ACE2 molecules (in magenta and cyan, respectively) with the distance between the C-terminal ends of the ACE2s indicated. (B) The top view of the S trimer in complex with three monomeric ACE2 molecules (in magenta, blue and cyan, respectively) with the distances between the C-terminal ends of the ACE2s indicated. (C) Superposition of the structure of the full-length ACE2 (PDB ID:6M17) and the structure of S-3 ACE2 complex.

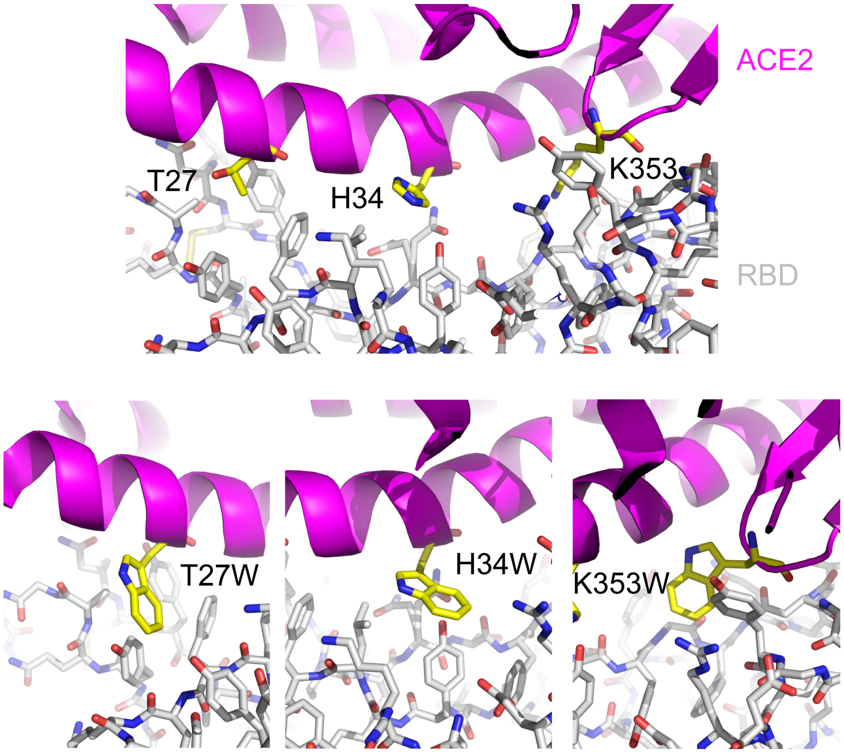


**Figure S5. Design of mutations at the ACE2-RBD interface.** Top, the interface between ACE2 in ribbon diagram in magenta and RBD in stick model from the complex structure (PDB ID: 6M0J) with T27, H34 and K353 from ACE2 indicated. Bottom, modeled T27W, H34W and K353W mutations that may enhance the hydrophobic interactions between ACE2 and RBD.

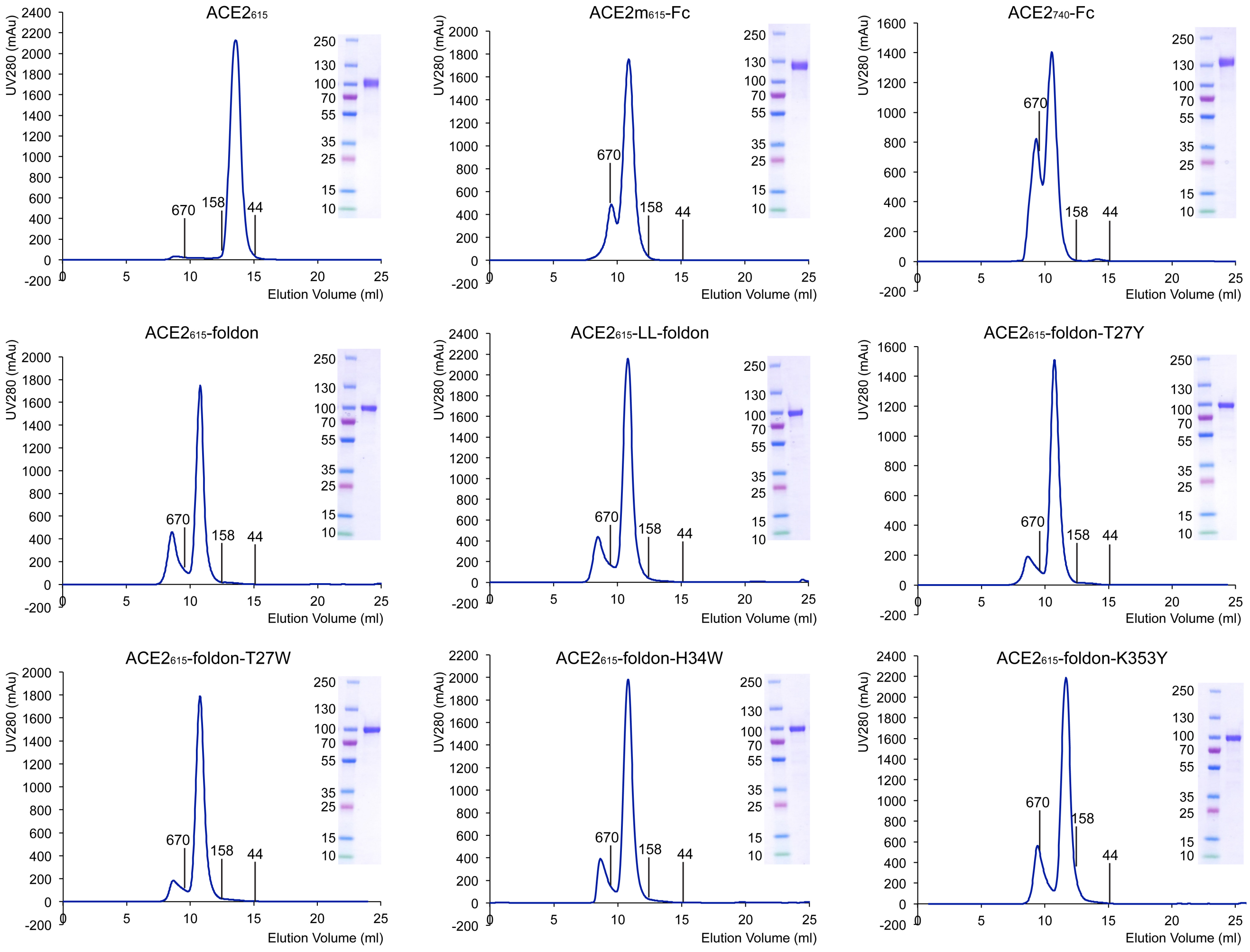


**Figure S6. Purification of ACE2 variants.** The purified ACE2 proteins were resolved by gel-filtration chromatography on a Superdex 200 column. The molecular weight standards include thyoglobulin (670 kDa), γ-globulin (158 kDa) and ovalbumin (44 kDa). Inset, peak fractions were analyzed by Coomassie stained SDS-PAGE.

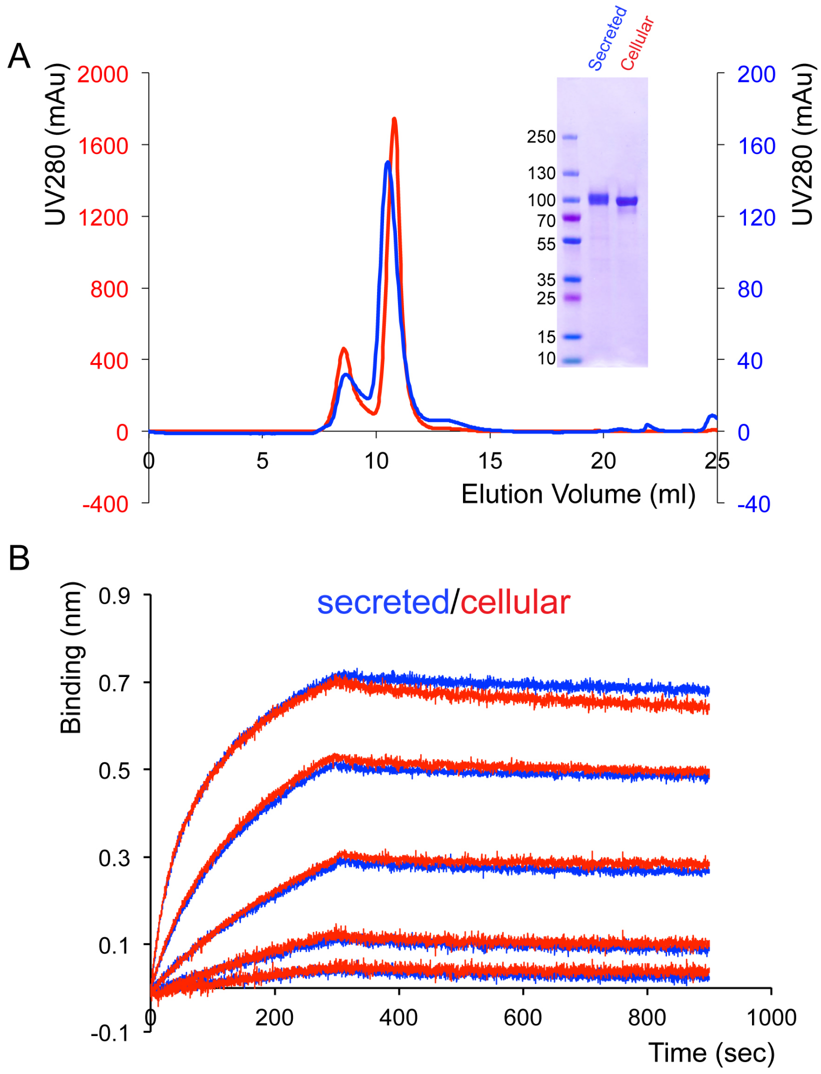


**Figure S7. Comparison of secreted and cellular ACE2$_{615}$-foldon protein.** (A) The purified ACE2$_{615}$-foldon protein either from cell supernatants (secreted) or cell lysates (cellular) was resolved by gel-filtration chromatography on a Superdex 200 column. Inset, peak fractions were analyzed by Coomassie stained SDS-PAGE. (B) Binding of ACE2$_{615}$-foldon to the stabilized soluble S trimer by bio-layer interferometry (BLI). The S protein was immobilized and subsequently dipped into the wells containing either secreted or cellular ACE2$_{615}$-foldon at various concentrations (0.617-50 nM). The sensorgrams for the secreted ACE2$_{615}$-foldon are in blue and those for the cellular ACE2$_{615}$-foldon in red.

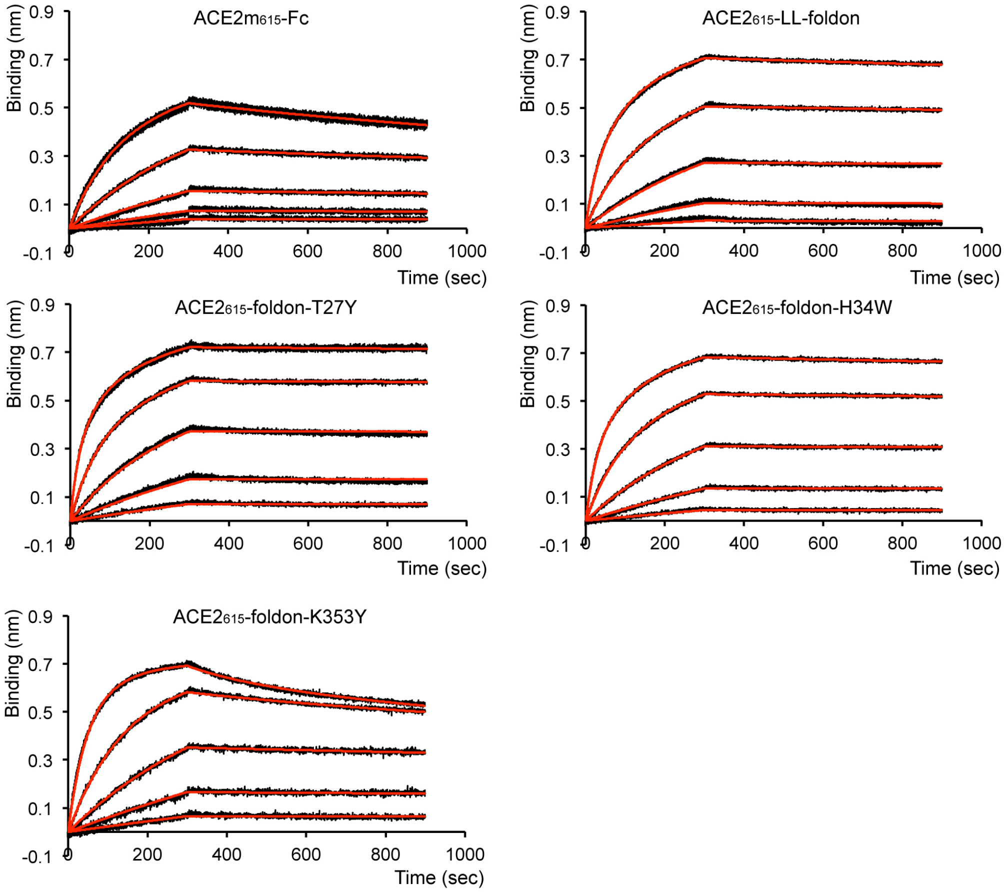


| Construct | KD (M) | KD Error | ka (1/Ms) | ka2 | ka Error | ka2 Error | kdis (1/s) | kdis2 | kdis Error | kdis2 Error |
|---|---|---|---|---|---|---|---|---|---|---|
| ACE2$_{615}$ | 7.68E-08 | 8.49E-10 | 1.10E+05 | | 1.09E+03 | | 8.44E-03 | | 4.17E-05 | |
| ACE2m$_{615}$-Fc | 2.23E-08 | 2.17E-09 | 3.12E+04 | 1.47E-01 | 5.94E+02 | 1.91E-01 | 6.96E-04 | 1.34E-02 | 6.63E-05 | 1.55E-02 |
| ACE2$_{740}$-Fc | 1.24E-08 | 5.68E-10 | 3.91E+04 | 6.50E-02 | 5.55E+02 | 1.94E-02 | 4.82E-04 | 9.01E-03 | 2.11E-05 | 2.13E-03 |
| ACE2$_{615}$-foldon | 1.15E-09 | 4.64E-11 | 1.46E+05 | 5.48E+00 | 2.61E+03 | 1.08E-01 | 1.68E-04 | 5.28E-01 | 6.08E-06 | 4.58E-02 |
| ACE2$_{615}$-LL-foldon | 6.24E-10 | 2.89E-11 | 1.33E+05 | 3.30E+00 | 2.54E+03 | 9.83E+00 | 8.26E-05 | 3.65E-01 | 3.49E-06 | 1.09E+00 |
| ACE2$_{615}$-foldon-T27Y | 9.45E-11 | 1.10E-11 | 2.09E+05 | 5.16E+00 | 3.12E+03 | 5.22E+00 | 1.97E-05 | 3.03E-01 | 2.27E-06 | 3.05E-01 |
| ACE2$_{615}$-foldon-T27W | 6.16E-11 | 6.54E-12 | 2.34E+05 | 2.37E-01 | 2.07E+03 | 2.07E-02 | 1.44E-05 | 2.29E-02 | 1.53E-06 | 2.12E-03 |
| ACE2$_{615}$-foldon-H34W | 3.66E-10 | 1.47E-11 | 1.58E+05 | 4.36E-01 | 2.26E+03 | 6.66E-02 | 5.80E-05 | 3.08E-02 | 2.18E-06 | 4.17E-03 |
| ACE2$_{615}$-foldon-K353Y | 3.23E-08 | 1.77E-09 | 9.96E+04 | 3.97E-01 | 5.16E+02 | 1.60E-01 | 3.22E-03 | 5.45E-03 | 1.75E-04 | 1.92E-03 |

**Figure S8. Binding of ACE2 variants to the stabilized soluble S trimer by bio-layer interferometry (BLI)**. The S protein was immobilized and subsequently dipped into the wells containing ACE2 proteins at various concentrations (0.926-75 nM for ACE2$_{615}$-Fc, 0.617-50 nM for the ACE2$_{615}$-foldon variants). Binding kinetics was evaluated using a bivalent model for all oligomeric ACE2s. The sensorgrams are in black and the fits in red. All the experiments were repeated at least twice with essentially identical results. Binding constants derived from the BLI experiments were also summarized.

**Table S1. EM data collection and reconstruction statistics**

| | | | | |
|---|---|---|---|---|
| Protein | Complex of ACE2 and SARS-CoV-2 S protein | | | |
| EMDB | | | | |
| Microscope | FEI Titan Krios | | | |
| Voltage(kV) | 300 | | | |
| Detector | Gatan K3 | | | |
| Magnification(nomina) | 105,000 | | | |
| Pixel size (Å/pix) | 0.825 | | | |
| Flux ($e^-$/pix/sec) | 14.83 | | | |
| Frames per exposure | 50 | | | |
| Exposure ($e^-/Å^2$) | 50.05 | | | |
| Dose per frame( $e/Å^2$) | 1.001 | | | |
| Defocus range (μm) | 1.6-2.7 | | | |
| Micrographs collected | 4,292 | | | |
| Particles extracted/final | 407,761/88,462 | | | |
| Symmetry imposed | C1 | | | |
| Class | No ACE2 | 1 ACE2 | 2 ACE2 | 3ACE2 |
| Map sharpening B-factor | -78.95 | -83.01 | -86.05 | -74.38 |
| Resolution at 0.143 FSC (Å) | 3.60 | 3.60 | 3.73 | 3.44 |
| **Model refinement and validation statistics** | | | | |
| PDB | | | | |
| Composition | | | | |
| Amino acids | 2906 | 3499 | 4106 | 4719 |
| Glycans | 59 | 70 | 80 | 90 |
| RMSD bonds (Å) | 0.010 | 0.006 | 0.013 | 0.010 |
| RMSD angles (◦) | 1.033 | 0.935 | 1.581 | 1.252 |
| Mean B-factors | | | | |
| Amino acids | 75.61 | 32.03 | 33.50 | 30.58 |
| Glycans | 131.78 | 51.75 | 55.17 | 47.49 |
| Ramachandran | | | | |
| Favored (%) | 90.76 | 93.76 | 90.60 | 91.80 |
| Allowed(%) | 9.20 | 6.22 | 9.18 | 8.18 |
| Outliers(%) | 0.04 | 0.03 | 0.22 | 0.02 |
| Rotamer outliers (%) | 0.31 | 0.07 | 10.39 | 12.10 |
| Clash score | 26.97 | 21.42 | 32.91 | 33.88 |
| C-beta outliers (%) | 0.00 | 0.03 | 0.65 | 0.07 |
| CC (mask) | 0.79 | 0.73 | 0.68 | 0.69 |
| MolProbity score | 2.45 | 2.24 | 3.31 | 3.33 |
| EMRinger score | 1.26 | 2.22 | 1.85 | 1.38 |
